# An Automatized Workflow to Study Mechanistic Indicators for Driver Gene Prediction with Moonlight

**DOI:** 10.1101/2022.11.18.517066

**Authors:** Astrid Saksager, Mona Nourbakhsh, Nikola Tom, Xi Steven Chen, Antonio Colaprico, Catharina Olsen, Matteo Tiberti, Elena Papaleo

**Affiliations:** Cancer Systems Biology, Technical University of Denmark, Copenhagen, Denmark; Department of Public Health Sciences, University of Miami Miller School of Medicine, Miami, FL, 33136, USA; Sylvester Comprehensive Cancer Center, University of Miami Miller School of Medicine, Miami, FL, 33136, USA; Vrije Universiteit Brussel (VUB), Universitair Ziekenhuis Brussel (UZ Brussel), Clinical Sciences, research group Reproduction and Genetics, Center for Medical Genetics, Brussels, 1090, Belgium; Brussels Interuniversity Genomics High Throughput core (BRIGHTcore), VUB-ULB, Brussels 1090, Belgium; Interuniversity Institute of Bioinformatics in Brussels (IB)2, Brussels 1050, Belgium; Cancer Structural Biology, Danish Cancer Society Research Center, Copenhagen, Denmark

**Author notes:** These two authors equally contributed to the work (Astrid Saksager, Mona Nourbakhsh).

**Keywords:** Driver Genes, Driver Mutations, Basal-like, Breast Cancer, Oncogenes, Tumor Suppressors

## Abstract

Prediction of tumor suppressors and oncogenes, also called driver genes, is an essential step in understanding cancer development and discovering potential novel treatments. We recently proposed Moonlight as a bioinformatics framework to predict driver genes and analyze them in a system-biology-oriented manner based on -omics integration. Moonlight uses gene expression as a primary data source and combines it with patterns related to cancer hallmarks and regulatory networks to identify oncogenic mediators. Once the oncogenic mediators are identified, it is important to include extra levels of evidence, called mechanistic indicators, to identify driver genes and to link the observed changes in gene expression to the underlying alteration that promotes them. Such a mechanistic indicator could be for example a mutation in the regulatory regions for the candidate gene or mutations in the regulator itself. In this work, we developed new functionalities and release Moonlight2, to provide the user with the mutation-based mechanistic indicator to streamline the analyses of this second layer of evidence. The function analyzes mutation information in a cancer cohort to classify them into driver and passenger mutations. Moreover, the function estimates the potential effect of a mutation on the transcriptional, translational, or protein structure/function level. Those oncogenic mediators with at least one driver mutation are retained as the final set of driver genes. We applied Moonlight2 and the newly developed function to a case study on Basal-like breast cancer subtype using data from The Cancer Genome Atlas. We found six oncogenes (*SF3B4, EBNA1BP2, KRTCAP2, ZBTB8OS, RUNX2*, and *POLR2J*) and ten tumor suppressor genes (*KIF26B, NR5A2, ARHGAP25, EMCN, ARL15, PCOLCE, TPK1, TEK, KIR2DL4*, and *GMFG*) containing a driver mutation in their promoter region, possibly explaining their deregulation. The Moonlight2R source code is available at https://github.com/ELELAB/Moonlight2R.

## Introduction

Cancer is a well-known and widespread disease and can in many cases lead to premature death. In 2020 it is estimated that 19 million people were diagnosed with cancer and almost 10 million people died because of cancer [1]. It is today in many (especially developed) countries the number one cause of premature deaths.

At the molecular level, different hallmarks of cancer have been identified [2–4]]. They are related to the deregulation of certain cellular functions, including increased cell proliferation, evasion of cell death, invasion, or escape of immune response. Cancer driver genes, which play important roles in connection to cancer hallmarks, are altered due to the accumulation of genomic alterations. They are known as tumor-promoting (oncogenes, OGs) or tumor suppressor genes (TSGs) [[5]. Mutations that activate OGs or inactivate TSGs drive tumor progression. Cancer driver genes can vary in cancer (sub)types, making them elusive to discover and annotate in a specific way. Even tumors with the same tissue of origin can be associated with different driver genes, complicating tumor stratification, accurate diagnosis, and targeted treatments [[6]]. In addition, a new group of genes has emerged in the last decade, known as ‘dual role’ driver genes or ‘double agents’. For example, Shen et al. 2018 found that out of 12 cancer types, breast cancer had the second highest occurrence of dual role genes such as *ARHGEF12, CBFA2T3, CDKN1B, DDB2*, and *FOXA1* [7]. Dual role driver genes can exhibit both oncogenic and tumor suppressor patterns depending on the cellular context [7–9].

Advances in cancer genomics and sequencing provided a multitude of data on profiling cancer samples, including data on gene expression, mutations, methylation, etc. To cite an example, The Cancer Genome Atlas (TCGA) accounts for more than 20 000 adult tumors [10,11]. Moreover, the Genomic Data Commons (GDC) has been developed as a portal for deposition and access to different cancer-omics data [12]. These data provide a precious source for the investigation and prediction of driver genes and their classifications.

As stated above, efforts in the discovery of driver genes are important not only for a fundamental understanding of cancer mechanisms. They have applicative interest since they can be investigated as drug targets [13,14] or they can be used as biomarkers to distinguish subtypes [15], which can increase the precision of prognosis [16]. Moreover, the knowledge of their dual role can help identify the most suitable treatment for a patient.

Many tools have been proposed to identify driver genes [17] but not all of them focus on the classification in TSGs or OGs [17]. In 2020, we contributed to this challenge by developing Moonlight, and its accompanying Bioconductor package, MoonlightR [9] which takes the biological context, the gene function, and its regulatory network into account. Moonlight is not solely based on changes in gene expression because this might be a poor indicator for driver genes [18]. The Moonlight framework requires at least an additional layer of evidence to link the changes in expression and regulation to what has been defined as a ‘mechanistic indicator’ [9]. Mechanistic indicators should help to understand the underlying reasons for a driver pattern (named oncogenic mediator) identified by MoonlightR. The evidence used can for example be chromatin accessibility, copy number variations, DNA methylation, and mutations [9]. The integration of these data allows for covering both genetic and epigenetic alterations that explain the changes in gene expression. However, in the original version of MoonlightR the step of definition of mechanistic indicators is left to the user and no specific protocols are provided. To tackle this challenge, streamline the process and provide a proper workflow for the identification of mechanistic indicators we devised Moonlight2R. In details, we provided a solution to the identification of mechanistic indicators based on mutation data, along with its implementation in a set of functions to streamline and automate the analysis. The functions are released within a new version of the package, Moonlight2R (https://github.com/ELELAB/Moonlight2R).

## Design and implementation

### Overview of Moonlight

For the sake of clarity, we will here give a brief overview of Moonlight, as originally conceived (**Fig 1**). The tool is designed to predict driver genes based on differentially expressed genes (DEGs) and additional layers of mechanistic indicators described in the original publication [9]. The first step in the Moonlight pipeline is a functional enrichment analysis (FEA) which determines if any of Moonlight’s 101 cancer-related biological processes are overrepresented among the DEGs. The next step is a gene regulatory network analysis (GRN) which considers the network of genes that the DEGs are a part of through mutual information. Following GRN, the Moonlight pipeline diverges into two modes: a machine-learning approach and an expert-based approach. The next step is an upstream regulatory analysis (URA) which will in both modes calculate the effect of the DEGs on either user-specified biological processes (the expert-based approach) or in all of Moonlight’s 101 biological processes (the machine learning approach). The final step is a pattern recognition analysis (PRA) which identifies the oncogenic mediators and divides them into putative OGs and putative TSGs. In the expert-based approach, this is done using patterns of the effect of DEGs on two biological processes with opposite effects on cancer. In the machine learning approach, the prediction of oncogenic mediators is carried out using a random forest classifier.

**Fig 1.**
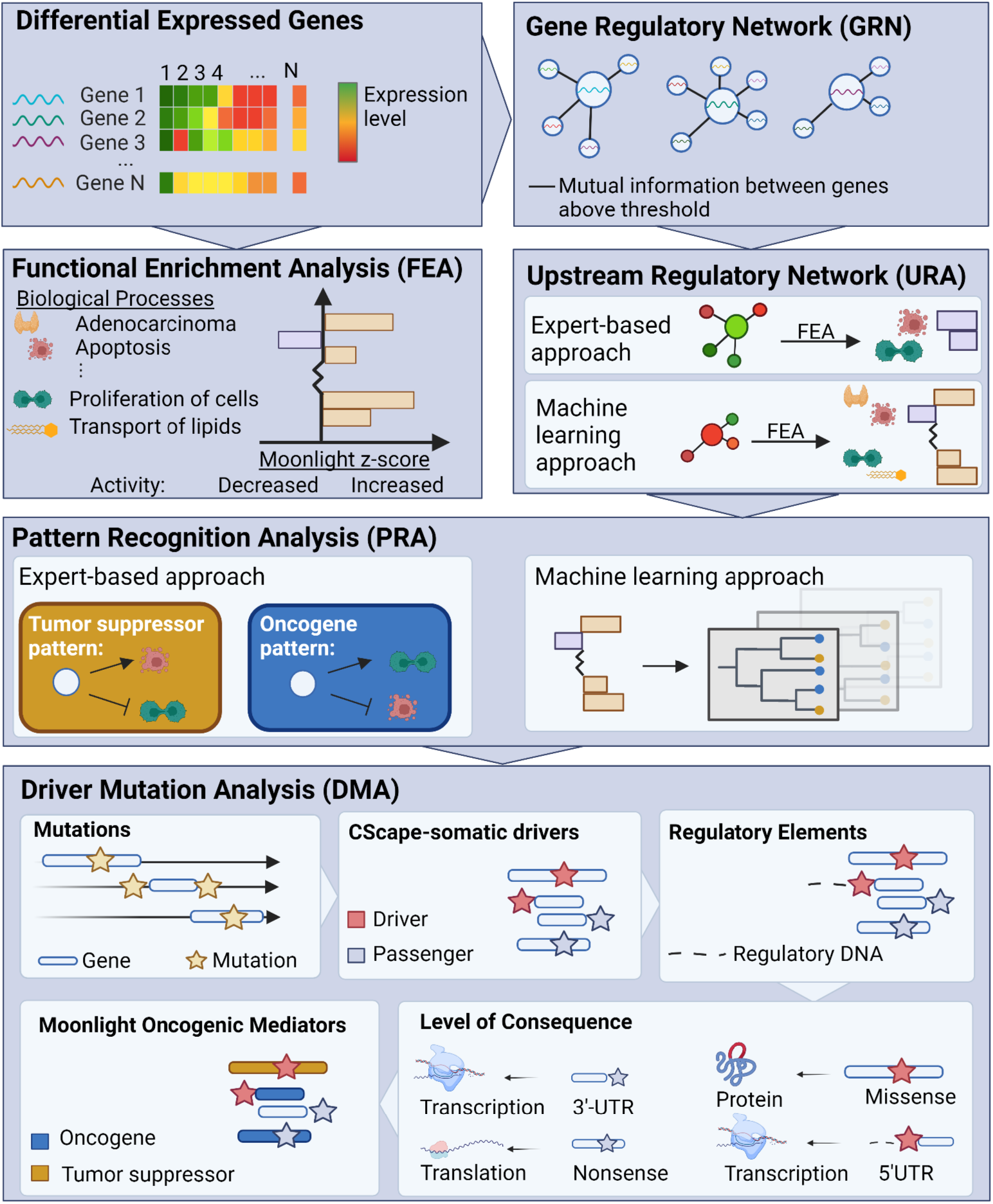
The Moonlight pipeline. The first step is a functional enrichment analysis (FEA) which determines if any of Moonlight’s 101 cancer-related biological processes are enriched among an input set of differentially expressed genes (DEGs). This is done through Fisher’s exact tests and Moonlight Process Z-scores. The Moonlight Process Z-scores indicate if the activity of the process is increased or decreased based on literature reportings and gene expression levels. The next step, a gene regulatory network analysis (GRN), creates gene networks for each DEG by calculating the mutual information between all pairs of DEGs. After GRN, the Moonlight pipeline diverges into an expert-based and a machine learning approach. The next step, an upstream regulatory analysis (URA), then evaluates the effect of each DEG on the biological processes through Moonlight Gene Z-scores. If the expert-based approach is selected, the Moonlight Gene Z-scores will only be calculated for chosen biological processes. If the machine learning approach is selected, the Moonlight Gene Z-scores will be calculated for all of Moonlight’s 101 biological processes. In the expert-based approach, pattern recognition analysis (PRA) identifies the oncogenic mediators which fit an OG or TSG pattern based on the Moonlight Gene Z-scores. The two chosen biological processes must have opposite effects on cancer (growing/blocking), e.g. proliferation of cells and apoptosis. In the machine learning approach, the prediction of oncogenic mediators is done using a random forest classifier. Following Moonlight’s primary layer, in Moonlight2, a secondary mutational layer is applied through the DMA step which identifies driver mutations. First, the mutations are divided into passengers and drivers by CScape-somatic. Then, regulatory elements from ENCODE are added. The consequence of the type of mutation is annotated to either the protein’s structure, the level of transcription or level of translation. Finally, the data is cross-referenced with the Moonlight driver genes.

### Design of the driver mutation analysis

In this paper, we present a new functionality to the Moonlight pipeline which allows for a mechanistic explanation of the predicted oncogenic mediators from Moonlight’s primary layer. The new function is called driver mutation analysis (DMA) and must be used subsequently to the PRA step. The function can distinguish between relevant and irrelevant mutations and help filter and prioritize the mutations found in a cohort of patients with the same cancer (sub)type. The function produces a summary of the oncogenic mediators and the assessed mutations, thereby strengthening the evidence for the Moonlight prediction of driver genes. The assessment by the function is visualized in **Fig 1** and has several internal steps. It removes all mutations from the Mutation Annotation Format (MAF) file that do not belong to any of the DEGs, and the remaining mutations are then classified as either drivers or passengers with CScape-somatic [19], then additional annotations are added on both mutation and gene levels. The details are provided in the next sections.

The function needs three inputs: the DEGs, the predicted oncogenic mediators, and a MAF file. The DMA function outputs the following: i) a list of oncogenic mediators with at least one driver mutation, now predicted as driver genes, ii) a table containing all annotations found to every DEG on both gene and mutational level, iii) a summary of the mutations found in the oncogenic mediators and finally, iv) a table containing the CScape-somatic file as if it was run outside of the DMA function.

CScape-somatic is a driver mutation predictor based on gradient-boosted decision trees. CScape-somatic defines a driver as a disease enabler that includes gain-of-function, loss-of-function, or both simultaneously [19]. CScape-somatic classifies somatic Single Nucleotide Polymorphisms (SNPs) on autosomes and scores each mutation with a number between zero and one, where one represents a highly likely driver mutation and zero represents a passenger mutation. We selected CScape-somatic for multiple reasons. First, it has the advantage of scoring mutations in both coding and noncoding regions of the genome. The possibility to cover driver variants in non-coding regions is central to the DMA function because the current Moonlight2 framework is still relying on gene expression data and thus are the mutations in the non-coding regions the most interesting to unveil as mechanistic indicators. Secondly, CScape-somatic provides data on mutations at a low computational cost since the machine-learning model has already scored the entire human genome and we do not need to include training/testing into our pipeline. In addition, CScape-somatic discriminate between germline neutral variants and somatic passenger mutations, which is not common to many available tools. We also considered the fact that the model has been trained on cancer samples and not any other disease, and as such should be more cancer-specific, which is an advantage according to a recent benchmark study [20]. The main limitation to consider is that it does not cover the X and Y chromosomes and that annotates only SNPs.

In our DMA workflow, we set the threshold to define a driver mutation for the CScape-somatic score to 0.5 (**Table 1**), as suggested in the original publication [19]. This means that all mutations with a score above 0.5 are denoted as driver mutations, and all mutations with a score less than 0.5 are denoted as passenger mutations. A threshold of 0.89 denotes driver mutations with high confidence, and so driver mutations with a score between 0.5 and 0.89 are labeled with low confidence. These thresholds and corresponding descriptions are retained in the output of DMA (see **Text S1** for more details).

**Table 1.** Thresholds of CScape-somatic scores annotating a mutation as a driver or passenger including the confidence of the mutation as a driver mutation.

| CScape-somatic score of mutation | Annotation of mutation by CScape-somatic | Confidence of mutation by CScape-somatic |
| --- | --- | --- |
| >0.89 (coding) | Driver | High confidence |
| >0.7 (non-coding) | Driver | High confidence |
| >0.5 | Driver | Low confidence |
| <0.5 | Passenger | Neutral |

### Predicting the level of consequence of mutations in DMA

The purpose of adding a mechanistic layer into Moonlight is to validate and explain the oncogenic patterns based on differential expression. For the mutations, it means that although a mutation might be a driver, it is not necessarily influencing the up- or downregulation of genes. We addressed this issue by categorizing mutations based on their position (e.g. 5’flank, 5’UTR, missense) and their type (e.g. SNP, insertion, deletion) into three categories which we call level of consequence: i) mutations which can influence the level (rate or amount) of transcription, ii) mutations which can influence the level of translation, and finally, iii) mutations which can influence a protein’s structure or function (see more details on the level of consequences in **Table S1** and **Text S2**). For instance, a mutation at the 5’flank of a gene can be inside a promoter. If the promoter is mutated, the transcription factor might bind stronger, weaker, or not at all, thereby causing a change in the expression level of the corresponding gene. On the other hand, a mutation at the translational start site (TSS) will not affect the transcription of the candidate gene but will influence the translation of the gene.

Besides annotating the predicted level of consequence of mutations, experimentally found promoter regions from the ENCODE consortium [21,22] are integrated in the DMA function. We downloaded the dataset from ENCODE (https://www.encodeproject.org/) with the following ENCODE identifier: ENCSR294YNI. If a mutation falls within a promoter region, we have added a column with the promoter start and end position to the output table containing a complete overview of all annotations belonging to the DEGs.

### Comparing driver genes with the Network of Cancer Genes

Finally, we incorporated data from the *Network of Cancer Genes* [[23] into the DMA function. From this, it is possible to cross-reference the oncogenic mediator annotation from Moonlight2 with findings from other cancer studies. In NCG, two categories of cancer genes are included. The first one contains known cancer driver genes with associated experimental support. The second one contains candidate cancer genes which somatic alterations are predicted to have a cancer driver role but without any experimental validation [23]. The Pubmed PMID identifier and the associated cancer types for each gene are also listed.

### Visualizing results of the DMA function

Additionally, two plotting functions have been added to the Moonlight package to visualize the results of the DMA function: *plotDMA()* which creates heatmaps of the driver classifications of mutations for the oncogenic mediators (Fig 2) and *plotMoonlight()* which visualizes the effect of genes on the biological processes calculated in the URA step (Fig 3).

**Fig 2.**
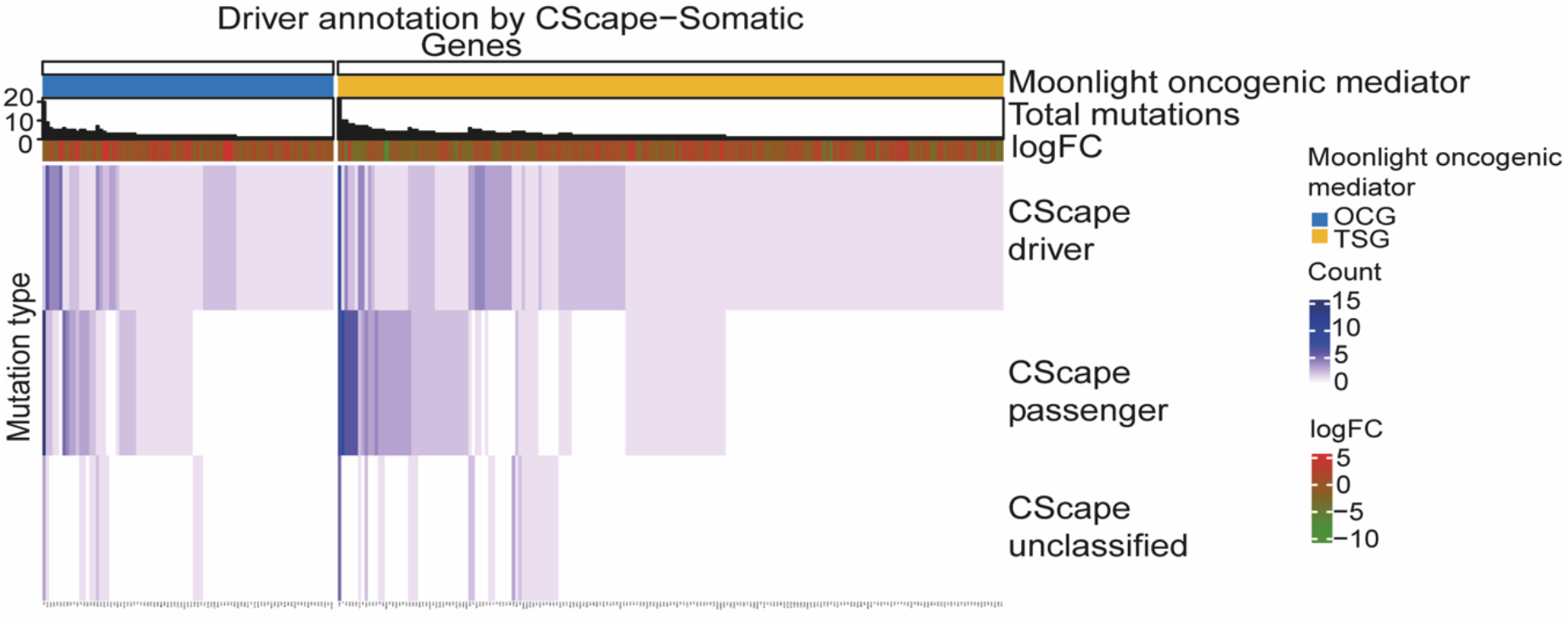
Classification of mutations in the 286 driver genes predicted by Moonlight. The 286 driver genes contain at least one driver mutation. Genes are in the columns while the mutation type classified by CScape-somatic is in the rows. The top blue block contains the predicted oncogenes (OGs) and the yellow block contains the predicted tumor suppressor genes (TSGs). The total number of mutations for the driver genes are shown in the top histogram. The red-green annotation bar represents the log2FC value of the driver genes. The purple lines in the heatmap indicate the number of driver, passenger, and unclassified mutations.

**Fig 3.**
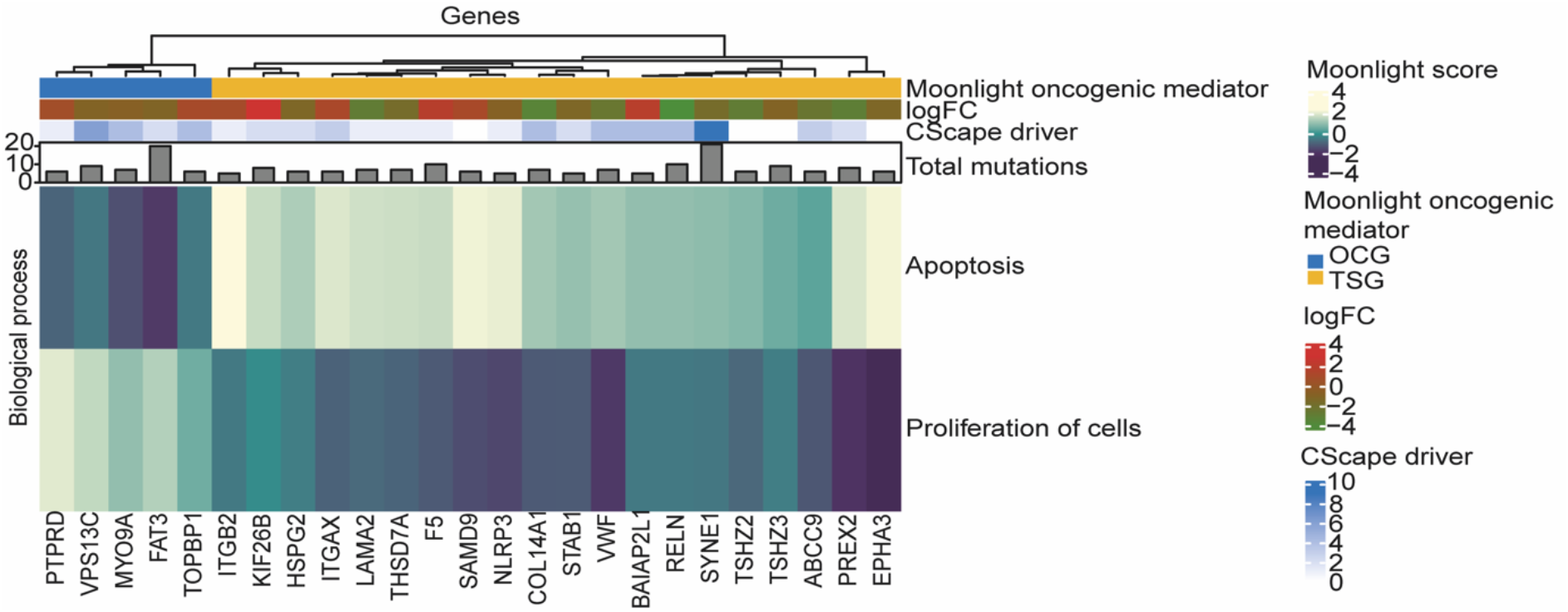
The top 25 oncogenic mediators with the highest total number of mutations. The rows indicate the genes and the columns refer to the Moonlight Gene Z-scores for the two biological processes selected in the expert-based approach: apoptosis and proliferation of cells. The top blue block contains the predicted oncogenes (OGs) and the yellow block contains the predicted tumor suppressor genes (TSGs). The blue-white annotation bar is the number of driver mutations. The red-green annotation bar represents the log2FC value of the oncogenic mediators. The gray bars represent the total number of mutations.

## Results

### Case study: Discovering driver genes in Basal-like breast cancer with Moonlight

To demonstrate the application of the new functionalities in MoonlightR, we conducted a case study on Basal-like breast cancer using data from TCGA. Gene expression, mutational and clinical data from the TCGA-BRCA project were retrieved via TCGAbiolinks [24,25] and further curated to include only the subtype Basal-like using the classification provided by the PanCancerAltas_subtype function of TCGAbiolinks [26] We performed a differential expression analysis (DEA) between the Basal-like subtype and normal samples to generate the input to Moonlight2. We implemented the steps in the Moonlight2 framework as outlined above (**Text S3**). Finally, we used the resulting tables from the DMA to select genes and mutations of interest which we investigated in the literature and other databases such as COSMIC [27] and TRRUST [28]. We predicted the driver genes in the context of the two biological processes apoptosis and proliferation of cells as these are well-known cancer hallmarks, as did in previous publications with Moonlight [9,29]. We also performed an enrichment analysis of the driver genes with the R package EnrichR [30,31] using the databases GO Molecular Function 2021, GO Biological Process 2021, and KEGG 2021 Human (**Text S3**).

### Applying the Moonlight pipeline on Basal-like breast cancer

From the DEA, we identified 9,300 DEGs between Basal-like breast cancer and normal samples which we then input to Moonlight2’s primary layer. This resulted in the prediction of 884 oncogenic mediators divided into 274 putative OGs and 610 putative TSGs. We then applied Moonlight2’s secondary mutational layer, implemented in the DMA function presented here, to mechanistically explain the predicted DEGs. This resulted in the classification of 4,125 driver mutations, 4,543 passenger mutations, and 1557 unclassified mutations. On the gene level, we found that 286 oncogenic mediators contained at least one driver mutation, resulting in our final set of driver genes. Of these 286 driver genes, 87 and 199 were predicted as OGs and TSGs, respectively (**Table 2** and **Table S2**).

**Table 2.** The upper part of the table contains the number of genes in different categories while the lower part contains the number of mutations. These categories are DEGs, oncogenic mediators, driver genes, and driver genes with transcriptional mutation(s). The numbers in brackets contain the mutations for OGs and TSGs, respectively. Note that the numbers are identical for driver mutations for oncogenic mediators and driver genes as this is the basis on which driver genes are chosen.

|  | DEGs | Oncogenic mediators | Driver genes | Driver genes with transcriptional mutation(s) |
| --- | --- | --- | --- | --- |
| **Genes** |  |  |  |  |
| OGs | - | 274 | 87 | 29 |
| TSGs | - | 610 | 199 | 63 |
| Total | 9,300 | 884 | 286 | 92 |
| Genes without mutations [OG/TSG] | 4539 | 378 [124/254] | - | - |
| **Mutations [OG/TSG]** |  |  |  |  |
| Driver mutations | 4125 | 411 [131/280] | 411 [131/280] | 156 [52/104] |
| Passenger mutations | 4543 | 531 [160/371] | 251 [82/169] | 78 [21/57] |
| Unclassified mutations | 1557 | 134 [35/99] | 52 [16/36] | 23 [6/17] |
| Total | 10225 | 1076 | 714 | 257 |
| Abbreviations: DEGs, differentially expressed genes; OGs, oncogenes; TSGs, tumor suppressor genes. |  |  |  |  |

We visualized the classification of the mutations of the 286 driver genes in a heatmap with the function *piotDMA()* (**Fig 2**). From Fig 2, we can notice that if an oncogenic mediator has a driver mutation, it will in about half the cases also have a passenger mutation. Moreover, we observe that most drivers only carry one driver mutation. We find one OG and one TSG which have around 20 total associated mutations whereas the rest of the predicted driver genes have ten or less total mutations.

### Exploration of oncogenic mediators and driver genes in terms of driver and passenger mutations

First, we explored the oncogenic mediators with a high number of mutations. In Fig 3, the top 25 oncogenic mediators with the highest number of total mutations are visualized in a heatmap, created with *plotMoonlight()*. It is clear that not all highly mutated oncogenic mediators contain few or any driver mutations e.g., *SAMD9, FAT3* and *SYNE1*. This clearly showcases the need to assess the mutations individually because the total number of mutations does not necessarily reflect driver status of the gene.

CScape-somatic has defined driver mutations as ‘disease-enablers’ [19], however, not all of them are placed such that they can directly interfere with the gene expression level. We therefore decided to focus on mutations with a transcriptional level of consequence. We found 92 driver genes containing in total 156 driver mutations which potentially affect the transcriptional level (**Table 2**). Of these transcriptional driver mutations, mutated promoter regions are of special interest because mutations in these sites can alter the transcription factor binding and thereby change the transcription level (**Fig 4** and **Table 3**). Of the 92 predicted Basal-like driver genes that had a driver mutation with a potential transcriptional level of consequence, we found that six OGs and ten TSGs had one driver mutation located within an experimentally validated promoter region from the ENCODE consortium [21,22]. We downloaded the dataset from ENCODE (https://www.encodeproject.org/) with the following ENCODE identifier: ENCSR294YNI. Of the driver mutations in these 16 genes (**Table 3**), *POLR2J*’s mutation scored the highest, suggesting high confidence of this driver mutation. These specific mutations are candidates for further structural studies and experimental investigation.

**Fig 4.**
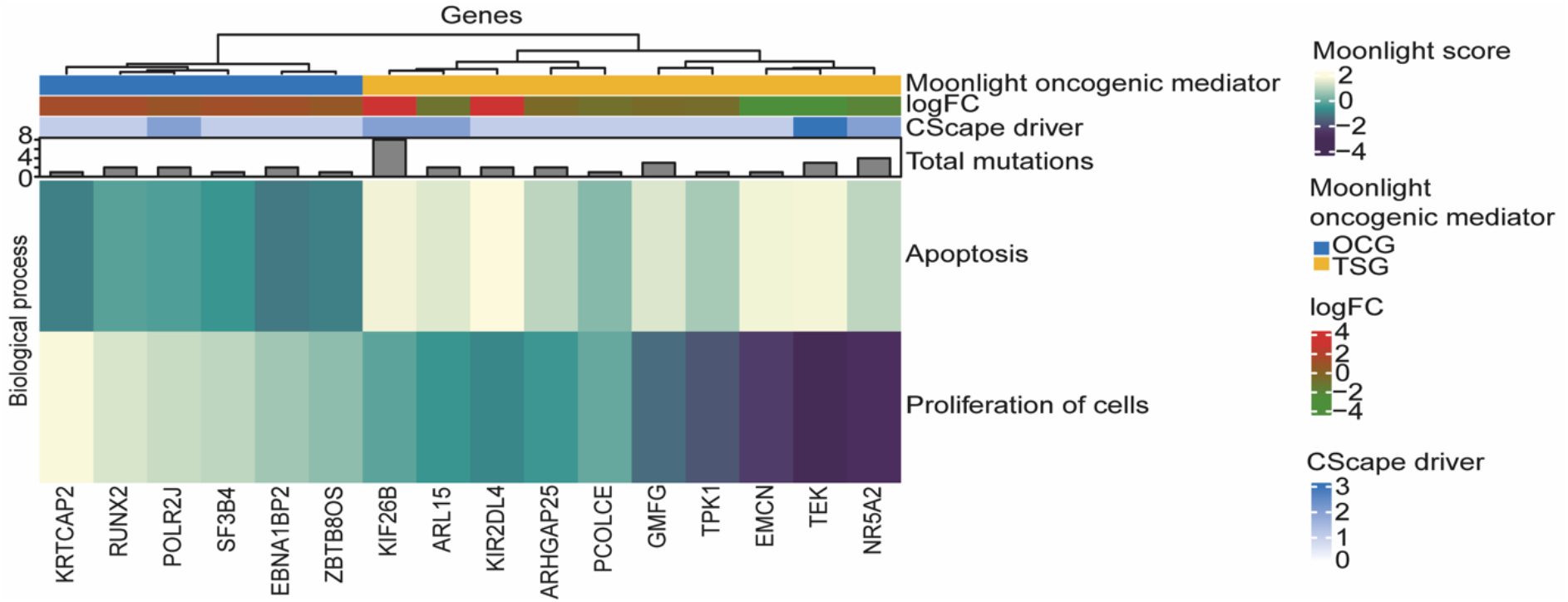
Oncogenes and tumor suppressors with a driver mutation in an experimentally validated promoter region. Six oncogenes (OGs) and ten tumor suppressors (TSGs) have a driver mutation located within an experimentally validated promoter region from ENCODE. The rows are genes, and the columns are the Moonlight Gene Z-scores for the two biological processes selected in the expert-based approach: apoptosis and proliferation of cells. The top blue block contains the predicted OGs and the yellow block contains the predicted TSGs. The blue-white annotation bar is the number of driver mutations. The red-green annotation bar represents the log2FC value of the oncogenic mediators. The gray bars represent the total number of mutations.

**Table 3.** Summary of driver mutations which are located in an experimentally validated promoter region from ENCODE. These mutations can therefore potentially alter the transcription level of the genes. The table includes which chromosome (Chr.) the mutation is placed on, Moonlight2 prediction of driver type (TSG/OG), SNP position in the chromosome according to grch38, mutation type, CScape-somatic score of mutation, and log2FC value of gene. *The mutation found in *EBNA1BP2* is the only mutation occurring in the protein coding region predicted by CScape-somatic. All other mutations were predicted as non-coding driver mutations.

| Chr. | Gene | Driver type (Moonlight2) | SNP Position | Classification of mutations | CScape-somatic score | Log2FC |
| --- | --- | --- | --- | --- | --- | --- |
| chr1 | KIF26B | TSG | 245155300 | 5'UTR | 0.746 | 3.03 |
| chr1 | SF3B4 | OG | 149927847 | 5'UTR | 0.682 | 1.12 |
| chr1 | NR5A2 | TSG | 200027802 | 5'UTR | 0.594 | -2.10 |
| chr1 | EBNA1BP2 | OG | 43172123 | 5'UTR | 0.7* | 1.06 |
| chr1 | KRTCAP2 | OG | 155173311 | 5'UTR | 0.683 | 1.27 |
| chr1 | ZBTB8OS | OG | 32650570 | 5'UTR | 0.631 | 0.654 |
| chr2 | ARHGAP25 | TSG | 68735192 | 5'UTR | 0.589 | -0.536 |
| chr4 | EMCN | TSG | 100517922 | 5'UTR | 0.532 | -2.67 |
| chr5 | ARL15 | TSG | 54310486 | 5'UTR | 0.528 | -1.11 |
| chr6 | RUNX2 | OG | 45422530 | Intron | 0.734 | 1.26 |
| chr7 | POLR2J | OG | 102478861 | 5'UTR | 0.815 | 0.770 |
| chr7 | PCOLCE | TSG | 100603943 | Intron | 0.625 | -0.949 |
| chr7 | TPK1 | TSG | 144835605 | 5'UTR | 0.524 | -0.712 |
| chr9 | TEK | TSG | 27109485 | 5'UTR | 0.697 | -2.66 |
| chr19 | KIR2DL4 | TSG | 54803614 | 5'Flank | 0.542 | 2.98 |
| chr19 | GMFG | TSG | 39335532 | Splice_Site | 0.599 | -0.642 |
| Abbreviations: OG, oncogenes; TSG, tumor suppressor gene; log2FC, log2 fold change; chr, chromosome; SNP, single nucleotide polymorphisms; UTR, untranslated region. |  |  |  |  |  |  |

On the flip side of the regulatory mechanisms lie the transcription factors (TFs) which bind to promoter regions and regulate transcription levels. Thus, we next sought to investigate the presence of any TFs in the 286 predicted Basal-like driver genes using the TRRUST database [28]. We found five and seven OGs and TSGs, respectively, which are TFs with a known mode of regulation (**Table 4**). Transcription factors can be of particular interest because they, like mutated promoters, can alter the transcription level. Though in this case it can be necessary to investigate all levels of consequences. For instance, if a mutation is placed in the coding region, the TF could have altered binding ability to the promoter, thereby changing the target’s expression level. The TF could also have its own transcription lowered, indirectly causing lowered transcription of the target. Such patterns might show up through further investigation of TF-target interactions.

**Table 4.** Basal-like driver genes predicted by Moonlight2 that are annotated as TFs in the TRRUST database with a known mode of action. Five and seven predicted OGs and TSGs are annotated as TFs in TRRUST, respectively. The log2FC value, number of driver mutations including type of driver mutations and which genes the TF has been found to regulate are shown. For the targets, the letters in parenthesis indicate mode of regulation: A = activation and R = repression.

| Gene | Basal-like driver gene (Moonlight2) | log2FC | Number of driver mutations | Target including mode of regulation by TF |
| --- | --- | --- | --- | --- |
| *APC* | OG | -0.752 | 1 | *AKT1* (R), *AMHR2* (R), *DNMT1* (R), *MYC* (R), *NOS2* (A), *ODC1* (R), *PTGS2* (R), *SGK1* (R) |
| *ERG* | TSG | -2.29 | 1 | *ADAMTS1* (A), *CXCL8* (R), *CXCR4* (A), *ENG* (A), *EPB41L3* (R), *EPB41L4B* (A), *ERG* (A), *FGF2* (R), *ICAM1* (R), *ILK* (R), *PIM1* (A), *SPP1* (A), *TDRD1* (A,R), *VIM* (A), *VWF* (A), *WNT11* (A) |
| *ILF3* | OG | 0.569 | 2 | *ACRV1* (A), *BIRC5* (A), *HLA-DRA* (R), *IL2* (A), *PLAU* (A) |
| *KLF11* | OG | -0.722 | 2 | *INS* (A) |
| *MITF* | TSG | -1.40 | 1 | *ACP5* (A), *BCL2A1* (R), *BEST1* (A), *CTSK* (A), *DCT* (A), *FOS* (A), *GPNMB* (A), *HIF1A* (A), *KIT* (A), *OCA2* (A), *PPARGC1A* (A), *SERPINF1* (A), *TRAP* (A), *TRPM1* (A), *TYR* (A), *TYRP1* (A) |
| *NFAT5* | OG | -0.949 | 4 | *CCL2* (R) |
| *NR5A2* | TSG | -2.10 | 2 | *ABCG5* (A), *ABCG8* (A), *CETP* (A), *CYP11B1* (A), *HSD3B2* (A), *NR5A2* (A), *STAR* (A) |
| *PGR* | TSG | -5.18 | 1 | *BCL2* (A), *CYP19A1* (R), *DUSP1* (A), *E2F1* (A), *ERBB2* (R), *ESR1* (A), *FOXP3* (R), *HLTF* (A), *IL10* (R), *KLK4* (A), *PTGS2* (R), *RELA* (R) |
| *PRDM1* | TSG | 0.562 | 1 | *CIITA* (R), *GCSAM* (R), *LMO2* (R), *MYC* (R) |
| *RUNX2* | OG | 1.26 | 1 | *ELANE* (A), *IBSP* (A, R), *LGALS3* (A), *MMP13* (A), *MMP2* (A), *SNAI2* (A), *SNAI3* (A), *TWIST1* (A), *VEGFA* (A), *VEGFC* (A) |
| *TAL1* | TSG | -2.39 | 1 | *ALDH1A2* (A), *CD34* (R), *ERG* (A), *MYB* (A), *MYCN* (A), *NFKB1* (R) |
| *TCF4* | TSG | -1.23 | 1 | *ABCB1* (A), *CCND1* (A), *CLEC4C* (A), *CNTNAP2* (A), *HECA* (R), *MITF* (R), *MYC* (A), *NOTUM* (A), *NRXN1* (A), *PLD1* (A), *PTEN* (R), *VEGFA* (A), *VIM* (A) |
| Abbreviations: TFs, transcription factors; OGs, oncogenes; TSGs, tumor suppressor genes; log2FC, log2 fold change; A, activation; R, repression. |  |  |  |  |

### Similarities and differences between TSGs and OGs

In the literature, it has been reported that OGs and TSGs generally promote and limit cell growth, respectively [32,33]. We sought to investigate possible differences between the two predicted classes of driver genes through an enrichment analysis on the 199 predicted TSGs and 87 predicted OGs (**Fig 5**).

**Fig 5.**
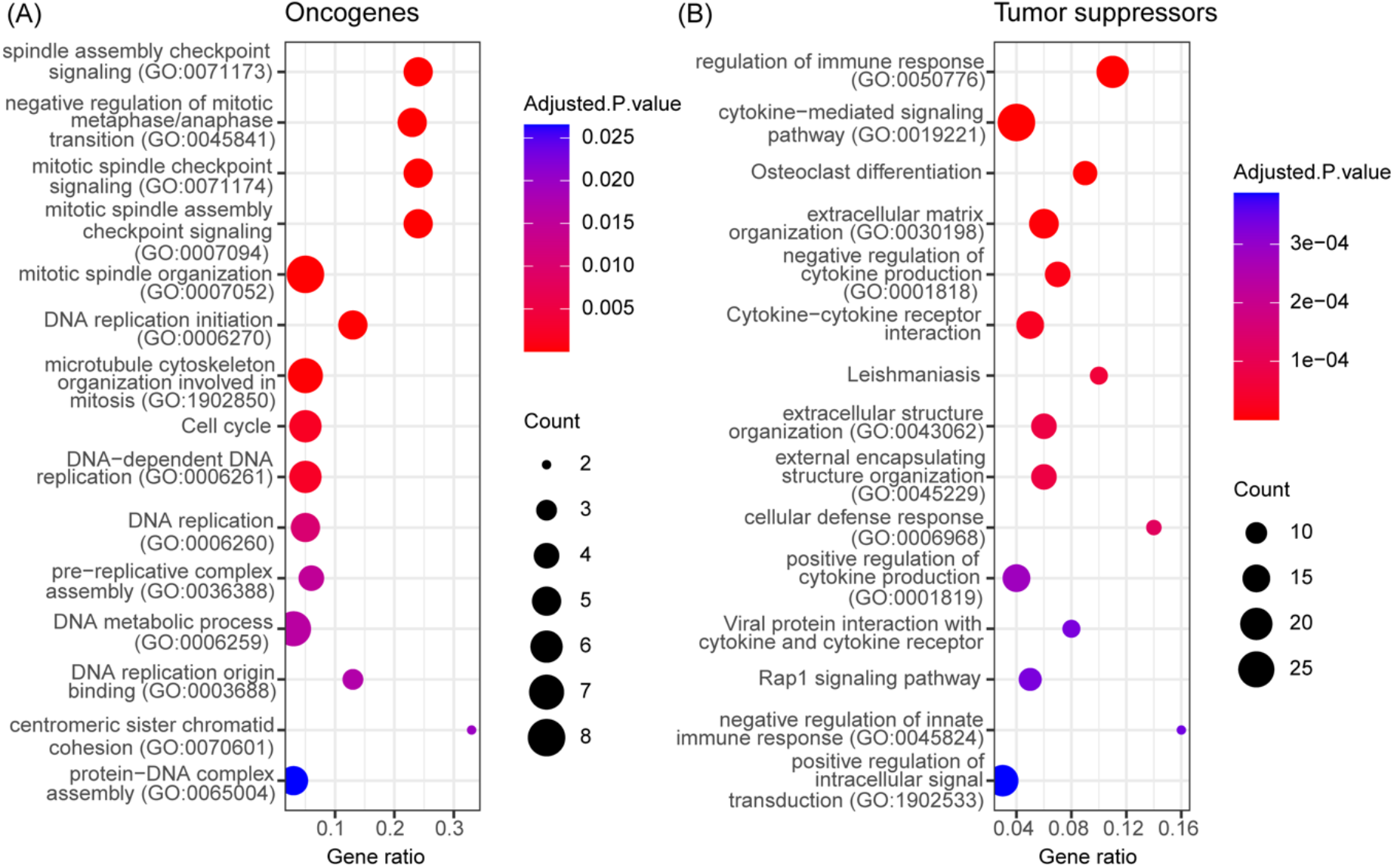
Enrichment analysis of 87 and 199 predicted oncogenes and tumor suppressors, respectively. The top fifteen most significantly enriched terms (adjusted p-value < 0.05) within the GO Molecular Function 2021, GO Biological Process 2021 and KEGG 2021 human databases among the (A) 87 predicted oncogenes (OGs) and (B) 199 predicted tumor suppressor genes (TSGs). The enriched terms are sorted on the adjusted p-value. The gene ratio refers to the ratio between the number of predicted TSGs or OGs intersecting with genes annotated in the given process and the total number of genes annotated in the given process. The colors of the points indicate the adjusted p-value and the sizes of the points represent the number of TSGs or OGs participating in the given process.

The enrichment analysis reveals expected oncogenic roles of the OGs. The top fifteen enriched terms among the OGs are related to DNA replication, spindle checkpoints, and cell cycle. Since the OGs are upregulated in Basal-like breast cancer compared to normal tissue, the activity of the enriched biological processes related to DNA replication and cell cycle is increased, potentially driving cancer progression. In contrast, the enriched terms among the TSGs are for the most part associated with immune response and regulation, with both positive and negative impacts, indicating immunological roles of the predicted TSGs. The enriched terms among the OGs and TSGs are related to the cancer hallmarks. For example, processes related to DNA replication and cell cycle are manifested in the *Sustaining proliferative signaling* hallmark.

Similarly, the immunological processes can play a role in the *Avoiding immune destruction* and *Tumor-promoting inflammation* hallmarks [2–4]. Collectively, these results indicate that Moonlight2, thanks to the integration of the core functions and the DMA step, is capable of finding two distinct classes of driver genes which both have strong associations to cancer related pathways. Moreover, only two enriched terms were shared between the TSGs and OGs when comparing the significantly enriched (adjusted p-value < 0.05), indicating that the two driver gene classes are highly distinct in terms of functional roles.

Besides investigating the functional profile of the predicted OGs and TSGs, we explored the difference between these two driver gene classes in terms of driver mutation types (**S1 Fig**). We noticed no distinct difference in the types of driver mutations between the OGs and TSGs, suggesting that it is not possible to classify the driver gene type based on driver mutation type alone.

### Comparison of predicted Basal-like driver genes with other findings and annotations

With the aim of investigating the novelty and consistency of Moonlight2’s driver gene annotations, we compared Moonlight2’s predictions with the Network of Cancer Genes (NCG) database (**Fig 6**). Most driver genes (i.e., 63 OGs and 133 TSGs) determined by Moonlight2 were not reported in the NCG database. The comparison between genes predicted as either OG or TSG with Moonlight2 and in NCG are listed in **Table S3.** The small overlap between Moonlight2’s predicted driver genes and NCG genes may also be attributed to NCG being a collection of genes across multiple cancer types and not only breast cancer. It could also be due to the lack of functional studies on some of the candidate genes with regards to their oncogenic or tumorsuppressor potential. Nevertheless, our predicted candidates would still need further investigation to support their driver signature and constitute an interesting dataset for the research community working with breast cancer subtypes.

**Fig 6.**
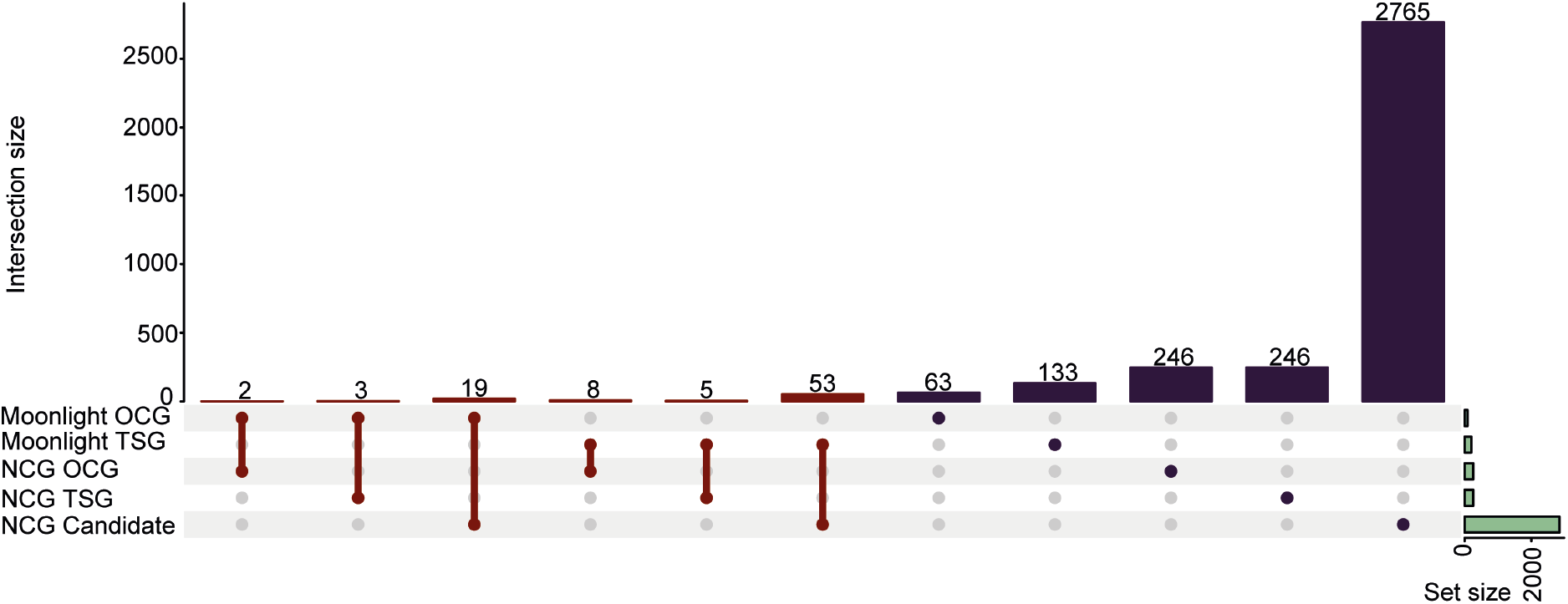
Comparison between Moonlight’s predicted driver genes and genes in the Network of Cancer Genes database. The red color represents genes in common between two groups and the purple color represents genes that are specific to one group. The green horizontal bar represents the total number of genes in the five sets.

One of the strengths of Moonlight is its classification of driver genes into TSGs and OGs which allows for the prediction of dual role genes - genes that are predicted as TSGs in one biological context but as OGs in another context [7,9]. We found 11 possible dual role driver genes which Moonlight2 has predicted as a TSG or an OG in the context of basal-like breast cancer, whereas NCG may have reported the gene functioning with the opposite driver gene type in possibly another cancer type. However, we found that none of the potential dual role genes had a reported cancer type by NCG. Thus, we could not establish the biological context of the dual role genes. Finally, we compared Moonlight’s basal-like driver genes with breast cancer genes reported in NCG to explore validity and consistency of our results. NCG contains 147 candidate driver genes associated with breast cancer, which are all marked as candidate driver genes. Comparing these 147 breast cancer genes with our 286 predicted driver genes revealed an overlap of five driver genes (*SYNE1, CTSS, PRKD1, STAB1* and *CD3G*) which are all predicted to be TSGs by Moonlight. Above, we also described *SYNE1* as the Moonlight2 predicted driver gene with the most driver mutations (**Fig 3**). Additionally, the four other genes have all been documented to be recurrently mutated across breast cancer cohorts [34–36]. Following the investigation of breast cancer related driver genes, we more specifically examined triple-negative breast cancer genes from NCG. 13 (CDKN2B, DNMT3B, EPHB1, IGF1R, MCL1, NOTCH3, PIK3CD, AURKA, DIS3, TBK1, SHQ1, RPTOR, and EPHA6) of the 147 breast cancer candidates in NCG are specifically associated with triple-negative breast cancer. These genes were also not identified by Moonlight2 as oncogenic mediators except for AURKA. Additionally, we only found five out of the 13 genes to be differentially expressed between Basal-like and normal samples used in our study, namely *DNMT3B, EPHB1, IGF1R, AURKA*, and *EPHA6*, meaning the other eight genes were not input to Moonlight, and consequently, we could not evaluate their driver gene potential. The fact that Moonlight2 with DMA did not predict any of these five genes as driver genes, suggesting that other mechanisms not dependent on changes in expression or mutations in non-coding regulatory elements could be at play, or features that are common only to a subset of patients and not identified in the TCGA dataset. To further investigate this direction, we retrieved the literature cited by NCG for each of the five genes and evaluated which was the expected driver alteration associated with them. DNMT3B, EPHB1, EPHA6 as for AURKA were annotated in NCG from the same study [37] but the results seem to mostly rely on predictions of sparse missense mutations or splice site and not clear experimental validation of the tumorigenic potential of these variants.

Few other tools besides Moonlight can classify driver genes as TSGs and OGs. We compared Moonlight basal-like predicted driver genes with another driver gene prediction tool, namely GUST [38]. We selected GUST due to the availability of precomputed publicly available results on breast cancer (https://liliulab.shinyapps.io/gust/). These predictions from GUST were however not basal-like specific. Compared to our results, we found a very small overlap of predicted driver genes. Only six genes were predicted as driver genes with both Moonlight2 and GUST (**Table S4**). Further comparing these results with NCG reveals that *JAK1* has previously been reported as an OG, and that *CCDC178* and *PRKD1* have been reported as candidate driver genes.

## Availability and future directions

The new functionality of Moonlight2 called DMA presented here is available in GitHub (https://github.com/ELELAB/Moonlight2R). The input, data and code for the case study is also available on GitHub (https://github.com/ELELAB/Moonlight2_DMA_basal_like). Once the Moonlight2R R package is loaded, example data of input and output files for all functions in Moonlight2 are provided.

In this study, we have mainly focused on driver mutations located in the promoter region as these mutations are the ones that most likely can explain the observed patterns of deregulated expression. Nevertheless, other types of mutations in both the coding and non-coding regions of the driver genes are of interest. For instance, missense mutations in genes involved with mRNA degradation could also be essential targets for further studies. Moreover, we envision complementing the promoter annotations by including other regions such as silencers and enhancers. These, along with a support to mutations occurring in transcription factors that are known to regulate with an oncogenic mediator predicted by Moonlight2 will be part of future developments. Additionally, we envision an automatized inclusion of additional -omics layers such as DNA methylation, copy number variation, and chromatin accessibility in future releases of Moonlight2.

## Acknowledgments

EP group’s research has been supported by Interregional Childhood Oncology Precision Medicine Exploration (iCOPE), a cross-Oresund collaboration between University Hospital Copenhagen, Rigshospitalet, Lund University, Region Skåne, and Technical University Denmark (DTU), Hartmanns Fond (R241-A33877) and Danmarks Grundforskningsfond (DNRF125). AC and XC were supported by grants from NCI R01CA200987, R01CA158472, and U24CA210954.

The results published here are in whole or in part based upon data generated by the TCGA Research Network: https://www.cancer.gov/tcga.

## Supporting information

**S1 Text. Design and implementation of Driver Mutation Analysis functionalities for Moonlight2R.**

**S2 Text. Details on definition of level of consequences.**

**S3 Text. Methods used in case study applying Moonlight2 on Basal-like breast cancer.**

**S1 Table. Level of consequence containing the predicted effect of different types of mutations on the transcription, translation, and protein structure/function level.**

**S2 Table. List of the 87 oncogenes and 199 tumor suppressors predicted in Basal-like breast cancer by Moonlight.**

**S3 Table. Overlapping driver genes between Moonlight2 predicted driver genes (TSG/OG) and driver genes reported in the NCG (TSG/OG).**

**S4 Table. Basal-like driver genes predicted by Moonlight2 that are annotated as TFs in the TRRUST database with a known mode of action.** Five and seven predicted OGs and TSGs are annotated as TFs in TRRUST, respectively. The log2FC value, number of driver mutations including type of driver mutations and which genes the TF has been found to regulate are shown. For the targets, the letters in parenthesis indicate mode of regulation: A = activation and R = repression.

**S1 Fig. Distribution of driver mutations.** Number of driver mutations belonging to 11 different mutation types of (A) The percentage out of the 131 driver mutations within 87 oncogenes (OGs) and (B) The percentage out of the 280 driver mutations within 199 tumor suppressor genes (TSGs) predicted in Basal-like breast cancer in percentages. The mutation types are classified in the MAF file of the TCGA-BRCA project.

**S2 Fig. Mutations per sample.** The sample is colored red if their mutation counts are outliers compared to average on the 99-percentile before filtering. (A) Mutations per sample before filtering of low-quality mutations and (B) mutations per patient after filtering of low-quality mutations.

## References

1. Sung H, Ferlay J, Siegel RL, et al. Global Cancer Statistics 2020: GLOBOCAN Estimates of Incidence and Mortality Worldwide for 36 Cancers in 185 Countries. CA Cancer J Clin 2021; 71:209–249

2. Hanahan D, Weinberg RA. The hallmarks of cancer. Cell 2000; 100:57–70

3. Hanahan D, Weinberg RA. Hallmarks of cancer: the next generation. Cell 2011; 144:646–674

4. Hanahan D. Hallmarks of Cancer: New Dimensions. Cancer Discov 2022; 12:31–46

5. Vogelstein B, Papadopoulos N, Velculescu VE, et al. Cancer Genome Landscapes. Science (1979) 2013; 339:1546–1558

6. Porta-Pardo E, Valencia A, Godzik A. Understanding oncogenicity of cancer driver genes and mutations in the cancer genomics era. FEBS Lett 2020; 594:4233–4246

7. Shen L, Shi Q, Wang W. Double agents: Genes with both oncogenic and tumor-suppressor functions. Oncogenesis 2018; 7:

8. Stepanenko AA, Vassetzky YS, Kavsan VM. Antagonistic functional duality of cancer genes. Gene 2013; 529:199–207

9. Colaprico A, Olsen C, Bailey MH, et al. Interpreting pathways to discover cancer driver genes with Moonlight. Nat Commun 2020; 11:

10. Hutter C, Zenklusen JC. The Cancer Genome Atlas: Creating Lasting Value beyond Its Data. Cell 2018; 173:283–285

11. Ganini C, Amelio I, Bertolo R, et al. Global mapping of cancers: The Cancer Genome Atlas and beyond. Mol Oncol 2021; 15:2823–2840

12. Grossman RL, Heath AP, Ferretti V, et al. Toward a Shared Vision for Cancer Genomic Data. New England Journal of Medicine 2016; 375:1109–1112

13. Liu Y, Hu X, Han C, et al. Targeting tumor suppressor genes for cancer therapy. Bioessays 2015; 37:1277–1286

14. Pagliarini R, Shao W, Sellers WR. Oncogene addiction: pathways of therapeutic response, resistance, and road maps toward a cure. EMBO Rep 2015; 16:280–296

15. Zhao L, Lee VHF, Ng MK, et al. Molecular subtyping of cancer: Current status and moving toward clinical applications. Brief Bioinform 2019; 20:572–584

16. Jackson SE, Chester JD. Personalised cancer medicine. Int J Cancer 2015; 137:262–266

17. Shi X, Teng H, Shi L, et al. Comprehensive evaluation of computational methods for predicting cancer driver genes. Brief Bioinform 2022; 23:

18. Pham VVH, Liu L, Bracken C, et al. Computational methods for cancer driver discovery: A survey. Theranostics 2021; 11:5553–5568

19. Rogers MF, Gaunt TR, Campbell C. CScape-somatic: Distinguishing driver and passenger point mutations in the cancer genome. Bioinformatics 2020; 36:3637–3644

20. Chen H, Li J, Wang Y, et al. Comprehensive assessment of computational algorithms in predicting cancer driver mutations. Genome Biol 2020; 21:

21. Luo Y, Hitz BC, Gabdank I, et al. New developments on the Encyclopedia of DNA Elements (ENCODE) data portal. Nucleic Acids Res 2020; 48:D882–D889

22. Dunham I, Kundaje A, Aldred SF, et al. An integrated encyclopedia of DNA elements in the human genome. Nature 2012; 489:57–74

23. Repana D, Nulsen J, Dressler L, et al. The Network of Cancer Genes (NCG): a comprehensive catalogue of known and candidate cancer genes from cancer sequencing screens. Genome Biol 2019; 20:

24. Colaprico A, Silva TC, Olsen C, et al. TCGAbiolinks: an R/Bioconductor package for integrative analysis of TCGA data. Nucleic Acids Res 2016; 44:e71

25. Silva TC, Colaprico A, Olsen C, et al. TCGA Workflow: Analyze cancer genomics and epigenomics data using Bioconductor packages. F1000Res 2016; 5:

26. Mounir M, Lucchetta M, Silva TC, et al. New functionalities in the TCGAbiolinks package for the study and integration of cancer data from GDC and GTEx. PLoS Comput Biol 2019; 15:e1006701

27. Tate JG, Bamford S, Jubb HC, et al. COSMIC: The Catalogue Of Somatic Mutations In Cancer. Nucleic Acids Res 2019; 47:D941–D947

28. Han H, Cho JW, Lee S, et al. TRRUST v2: an expanded reference database of human and mouse transcriptional regulatory interactions. Nucleic Acids Res 2018; 46:D380–D386

29. Lucchetta M, da Piedade I, Mounir M, et al. Distinct signatures of lung cancer types: Aberrant mucin O-glycosylation and compromised immune response. BMC Cancer 2019; 19:

30. Chen EY, Tan CM, Kou Y, et al. Enrichr: interactive and collaborative HTML5 gene list enrichment analysis tool. BMC Bioinformatics 2013; 14:

31. Xie Z, Bailey A, Kuleshov M v., et al. Gene Set Knowledge Discovery with Enrichr. Curr Protoc 2021; 1:

32. Croce CM. Oncogenes and cancer. N Engl J Med 2008; 358:502–511

33. Wang LH, Wu CF, Rajasekaran N, et al. Loss of Tumor Suppressor Gene Function in Human Cancer: An Overview. Cellular Physiology and Biochemistry 2018; 51:2647–2693

34. Ciriello G, Gatza ML, Beck AH, et al. Comprehensive Molecular Portraits of Invasive Lobular Breast Cancer. Cell 2015; 163:506–519

35. Encinas G, Sabelnykova VY, de Lyra EC, et al. Somatic mutations in early onset luminal breast cancer. Oncotarget 2018; 9:22460–22479

36. Berger AC, Korkut A, Kanchi RS, et al. A Comprehensive Pan-Cancer Molecular Study of Gynecologic and Breast Cancers. Cancer Cell 2018; 33:690–705.e9

37. Weisman PS, Ng CKY, Brogi E, et al. Genetic alterations of triple negative breast cancer by targeted next-generation sequencing and correlation with tumor morphology. Modern Pathology 2016; 29:476–488

38. Chandrashekar P, Ahmadinejad N, Wang J, et al. Somatic selection distinguishes oncogenes and tumor suppressor genes. Bioinformatics 2020; 36:1712–1717

