## Supplementary Text 1 for "An Automatized Workflow to Study Mechanistic Indicators for Driver Gene Prediction with Moonlight"

**S1 Text. Considerations and choices of Driver Mutation Analysis implementation.**

*Reasons for choosing a MAF file as input to DMA*

We chose a cohort-based file like MAF as input to DMA because Moonlight is generally built to provide cohort-level predictions rather than patients-specific predictions. Moreover, Moonlight is designed for easy integration with TCGA data which also contain publicly available MAF files. The alternative to MAF files is VCF files, which are patient-specific, however, they are often protected, which complicates obtaining such files. Besides, several tools exist which can produce a MAF file by combining several VCFs together. Hence, even if a user only has VCF files available, they can easily be converted to a MAF file.

*Runtime of DMA*

The runtime of DMA is largely dependent on the number of mutations in the MAF input, however, it scales linearly. The DMA function ran for approximately twenty minutes with one core for the Basal-like case study presented in this paper while the full TCGA-BRCA MAF file containing data for all four subtypes can be processed in one and a half hours. The CScape-somatic tables must be downloaded by the user from their website. These tables are quite large with the coding and non-coding files occupying 632 megabytes and ~46 gigabytes, respectively, and may therefore take a while to download. All output files of DMA are provided in rda format to reduce disk space.

*Inclusion of CScape-somatic driver mutation prediction tool in DMA*

The first and most essential part of DMA is the division of mutations into driver and passengers with CScape-somatic, a driver mutation predictor published by Rogers et al. in (2020) [1]. The authors developed the model based on integrative single kernel learning which estimated the likelihood of mutations being cancer drivers. In the work, they also tested several different models with forward selection of more than 30 features and evaluated them on balanced accuracy by leave-one-chromosome-out cross-validation. The training set containing true negative and true positive mutations was based on selected mutations from COSMIC. The final kernel consisted of gradient boosted decision trees. The noncoding model includes the following features determined from forward selection: *conservation*, *local mutation frequency*, *distances from gene features*, *sequence uniqueness* and *GC content*. The coding model includes the same features as the noncoding model with the additional inclusion of the features *functional elements* and *spectrum groups*. CScape-somatic has been specifically trained to distinguish between somatic drivers and somatic passengers, compared to the older version of CScape [2], which was based on neutral germline mutations. It is therefore vital that the mutational input to DMA is somatic.

One limitation to CScape-somatic is the fact that it does not cover the X and Y chromosomes. In this case study, all patients were female so none of the mutations could be placed on a Y chromosome. However, 15 oncogenic mediators predicted by Moonlight's primary layer were placed on the X chromosome and even though they all had mutations, we could not score these through CScape-somatic. Moreover, CScape-somatic only annotates SNPs which means that indels are not classified into drivers and passengers. Indels are a well-known type of genomic alteration causing cancer [3][ 10.1038/nm.4002]. For example, in our case study of Basal-like, we found 41 oncogenic mediators which only contained indels and were excluded from further analysis. A possible future solution could be implementing PredCID, a tool specifically designed to predict indels driver status in cancer [4].

*Details of installation, run and test data supplied in MoonlightR2*

The new software is available to download as an R package through the Github repository https://github.com/ELELAB/MoonlightR2/, where any future updates of Moonlight will also be deposited. Included in the package is test data to all functions and their subsequent results based on TCGA-LUAD. Further a vignette describes how this data is compiled, how the functions of Moonlight are run and how to visualize the results. All test data and tables included in the package are described and documentation can be called once the package is installed.

### References

1. Rogers MF, Gaunt TR, Campbell C. CScape-somatic: Distinguishing driver and passenger point mutations in the cancer genome. Bioinformatics 2020; 36:3637–3644
2. Rogers MF, Shihab HA, Gaunt TR, et al. CScape: A tool for predicting oncogenic single-point mutations in the cancer genome. Sci Rep 2017; 7:
3. Vogelstein B, Papadopoulos N, Velculescu VE, et al. Cancer genome landscapes. Science (1979) 2013; 340:1546–1558
4. Yue Z, Chu X, Xia J. PredCID: Prediction of driver frameshift indels in human cancer. Brief Bioinform 2021; 22:
