## Supplementary Text 2 for "An Automatized Workflow to Study Mechanistic Indicators for Driver Gene Prediction with Moonlight"

**S2 Text. Level of consequence.**

***<u>Level of consequence - introduction:</u>***

A mutation of a certain type and class can result in different effects in the central dogma. The CScape-somatic driver mutation prediction tool scores SNPs according to their cancer driver effect. However, it does not explicitly explain which part in the central dogma it affects. To counter this, we created three new tables assessing these effects. Each table contained the variant classification in the rows and the variant type in the columns (S1 Table). The values are binary values, reflecting if this category of mutation had the possibility of affecting either transcriptional levels, translational levels or the protein structure and/or function. These definitions only apply to the genes in which the mutation is located. We did not consider cases where for instance a protein disruptive mutation will affect the transcription level if it is in a transcription factor.

***Level of consequence - definitions:***

Transcription/level of transcript:

The possible change in level/amount of transcript can be caused either by regulatory changes in the transcription process or mechanisms surrounding the mRNAs stability and degradation.

1: The amount of mRNA can be affected

0: The amount of mRNA cannot be affected

Translation/level of peptide:

The possible change in level/amount of peptide can be caused by regulatory changes in the translation processes, mechanisms surrounding peptide stability and degradation or the subsequent effect of up- or downregulation of mRNA available to translate. The level will also be indirectly affected if the level of transcript is changed.

1: The amount of peptide can be affected

0: The amount of peptide cannot be affected

Protein structure or function:

The possible mutations which are contained in the final peptide after transcription and translation of the gene and can cause an altered structure or function of the protein.

1: The protein structure and/or function can be altered

0: The protein structure and/or function cannot be altered

***Variant types:***

This variant types are the ones found in a given Mutation Annotation Format (MAF) file from TCGA (https://docs.gdc.cancer.gov/Data/File_Formats/MAF_Format/#notes-about-gdc-maf-implementation) and are abbreviated as follows:

SNP: Single Nucleotide Polymorphism
INS: Insertion
DEL: Deletion
DNP: Di-nucleotide polymorphism, two consecutive nucleotides
TNP: Tri-nucleotide polymorphism, three consecutive nucleotides
ONP: Oligo-nucleotide polymorphism, four or more consecutive nucleotides
Consolidate: “Consolidation is used to indicate a site that was initially reported as a variant but subsequently removed from further analysis because it was consolidated into a new variant. For example, a SNP variant incorporated into a TNP variant.”

***Variant classification:***

The variant classifications are assigned by VEP as part of the GDC bioinformatics pipeline, DNA-seq analysis, in the somatic aggregation workflow. A variant is annotated if it is within 5kb of a feature (https://docs.gdc.cancer.gov/Data/Bioinformatics_Pipelines/DNA_Seq_Variant_Calling_Pipeline/#somatic-aggregation-workflow). The sequence ontology used by VEP is listed here: http://www.ensembl.org/info/genome/variation/prediction/predicted_data.html#consequences

***<u>Structure for how description of each mutation is provided below:</u>***

<u>*Definition:*</u> Definition of the mutation based on sequence ontology (SO).
<u>*Argument for the decision*</u>: This is an explanation on how/how not a mutation affects the different levels and a clarification if a variant type is not possible for this classification.
If there is a parenthesis around (1), it is because the variant does not directly alter the translational level, but the transcript levels are changed due to the mutation which could subsequently change the peptide levels.
<u>*Result*</u>:

1. Transcription
2. Translation
3. Protein structure/function

<u>*Example in literature:*</u> A reference supporting the most important arguments. When it is possible, the paper will be about a specific mutation variant which has been shown to affect one or more of the level of consequences and have been related to cancer. However, in some cases it has only been possible to find reviews describing the mutation variants and the genomic regions.

***Results:***

**5'Flank:**

*Definition*: SO: “A sequence variant located 5' of a gene”

*Argument*: A promoter is found upstream of the transcriptional start site of the gene and is therefore found in the 5’flank region. If a promoter is mutated, it could alter the binding ability of a transcription factor, thereby changing the gene expression. It is not in the coding region and will not affect the primary structure of mRNA and thereby not the protein.

*Result:*

Transcription: 1
Translation: (1)
Protein: 0

*Example in literature*: 10.1186/s12885-022-09178-z

**5'UTR:**

*Definition:* SO: “A UTR variant of the 5' UTR”

*Argument:* There are many elements in the 5’ untranslated region which can alter transcription levels such as uORFs. Other elements e.g. 5’cap structures or secondary structures can change the mRNA stability and degradation or the efficiency of translation. As the name indicates, the region is not translated and cannot affect the protein structure.

*Result:*

Transcription: 1
Translation: 1
Protein: 0

*Example in literature:* 10.1038/s41594-020-0465-x, 10.1038/s41467-021-24445-6

**TSS:**

*Definition:* SO: “A codon variant that changes at least one base of the canonical start codon”

*Argument:* This is the translational start site, which is placed at the beginning of the coding region. If the start codon is lost and there is no alternative start codon the protein will not be translated at all. If there is an alternative start codon the protein will be truncated.

*Result:*

Transcription: 0
Translation: 1
Protein: 1

*Example in literature:* 10.1016/j.ymthe.2019.11.022

**Missense**:

*Definition:* SO: “A sequence variant, that changes one or more bases, resulting in a different amino acid sequence but where the length is preserved”

*Argument:* Missense mutations can only be found in the coding region. The mutation can therefore change the protein structure, but will not affect any regulatory regions

that could alter transcription or translation. A missense mutation is per definition not an insertion or deletion.
*Result:*
Transcription: 0
Translation: 0
Protein: 1
*Example in literature:* 10.1038/s41436-018-0018-4

**Nonsense**:
*Definition:* SO: "A sequence variant whereby at least one base of a codon is changed, resulting in a premature stop codon, leading to a shortened transcript"
*Argument:* A nonsense mutation must per definition be part of the transcript. If a transcript contains a premature termination codon, a process called nonsense-mediated decay is activated which will break down the mRNA. This will therefore also cause downregulation of the peptide. If this process is not activated, the protein might have an altered structure.
*Result:*
Transcription: 1
Translation: (1)
Protein: 1
*Example in literature:* 10.1016/j.gde.2017.10.007

**Nonstop**:
*Definition:* SO: "A sequence variant where at least one base of the terminator codon (stop) is changed, resulting in an elongated transcript"
*Argument:* An elongated polypeptide will change the protein. It has been shown that some nonstop mutations will cause vast degradation of the protein via the ubiquitin–proteasome system. There seems to be no reduction of nonstop mutated mRNA, but the translation is significantly halted.
*Result:*
Transcription: 0
Translation: 1
Protein: 1
*Example in literature:* 10.1038/s41556-020-0551-7, 10.1038/sj.emboj.7601679

**Splice mutations:**
The result of a splice site or splice region is difficult to assess and is yet to be well understood. It could result in retention of introns, skipping of exons, introduction of new splice sites (these are, however, not included in these classifications) and change the balance of isoforms (10.1136/jmg.2004.029538).

**Splice Site:**
*Definition:* SO: Splice_acceptor_variant "splice variant that changes the 2 base region at the 3' end of an intron" or Splice_donor_variant "A splice variant that changes the 2 base region at the 5' end of an intron"
*Argument:* The literature is sparse on the effect of splice sites mutations. There are, however, examples in the literature of splice donor site mutations which cause complete lack of expression of the gene. It cannot be rejected that these mutations

cause variation on the transcription and translational level. Through retention of introns or skipping of exons, the protein would change, though possibly just to a different isoform.

*Result:*

Transcription: 1

Translation: 1

Protein: 1

*Example in literature:* 10.1016/j.ymeth.2017.06.001

**Splice Region:**

*Definition:* SO: “A sequence variant in which a change has occurred within the region of the splice site, either within 1-3 bases of the exon or 3-8 bases of the intron”

*Argument:* The literature is sparse on the effect of splice sites mutations. There are, however, examples in the literature of splice donor site mutations which cause complete lack of expression of the gene. It cannot be rejected that these mutations cause variation on the transcription and translational level. As the splice region can be placed within the exon, this would affect the protein structure. Even if the mutation were in the intron, the protein would change through retention of introns or skipping of exons, though possibly just to a different isoform.

*Result:*

Transcription: 1

Translation: 1

Protein: 1

*Example in literature:* See *splice site* section described above.

**Silent:**

*Definition:* SO: “A sequence variant where there is no resulting change to the encoded amino acid”

*Argument:* Per definition the silent mutations are only in the coding region. If they are found outside the coding region, they will be categorized by region. It has been shown that translation rates are, among other features, affected by codon-bias and secondary structure of mRNA, both which could be changed with a silent mutation. A silent mutation cannot be an insertion or a deletion.

*Result:*

Transcription: 0

Translation: 1

Protein: 0

*Example in literature:* 10.1146/annurev-biochem-071320-112701, 10.1016/j.molcel.2015.05.035

**Frameshift and inframe mutations:**

No frameshift ins/del or any in frame ins/dels can be DNP, TNP or ONP as these are not insertions or deletions but polymorphisms, exchange of one or more nucleotides with another. Frameshift or in-frame mutations will always at least partly cover an exome, the coding region. These mutations will not create nonsense or non-stop nor will they affect splice regions/sites as they would then have been classified as such. Therefore, they will only be within the exon and can therefore only affect the protein structure. Frameshift insertion and deletion share the same sequence ontology.

**In Frame Deletion:**

*Definition:* SO: "A sequence variant which causes a disruption of the translational reading frame, because the number of nucleotides inserted or deleted is not a multiple of three"
*Argument:* These mutations will shorten the peptide. In frame deletion mutations can only have the variant type DEL.
*Result:*
Transcription: 0
Translation: 0
Protein: 1
*Example in literature:* 13030/qt5t22m5gk

**In Frame Insertion:**

*Definition:* SO: "A sequence variant which causes a disruption of the translational reading frame, because the number of nucleotides inserted or deleted is not a multiple of three"
*Argument:* These mutations will lengthen the peptide. In-frame insertion mutations can only have the variant type INS.
*Result:*
Transcription: 0
Translation: 0
Protein: 1
*Example in literature:* 10.1038/s41598-019-44584-7

**Frameshift Deletion:**

*Definition:* SO: "An inframe non synonymous variant that deletes bases from the coding sequence"
*Argument:* These mutations will shorten the peptide. This can only have the variant type DEL.
*Result:*
Transcription: 0
Translation: 0
Protein: 1
*Example in literature:* 10.1007/s10549-016-4100-9

**Frameshift Insertion:**

*Definition:* SO: "An inframe non synonymous variant that inserts bases into in the coding sequence"
*Argument:* Frameshift insertion mutations can only have the variant type INS. These mutations will lengthen the peptide.
*Result:*
Transcription: 0
Translation: 0
Protein: 1
*Example in literature:* 10.1038/s41598-017-09287-x

**Intron:**

*Definition:* SO: “A transcript variant occurring within an intron“
*Argument:* Many examples of mutations in the deep intronic regions which alter or disrupt splicing of mRNA have been found. This can occur when an intronic enhancer or silencer are hit and will result in an altered peptide. Further, some promoters have been shown to interact with enhancers placed inside introns. As this could affect the binding ability of transcription factors, the transcription level could be altered.
*Result:*
Transcription: 1
Translation: 1
Protein: 1
*Example in literature:* 10.1007/s00439-017-1809-4

**3'UTR:**

*Definition:* SO: “A UTR variant of the 3' UTR”
*Argument:* The expression of genes is regulated by miRNA binding to untranslated regions (UTRs) at the 3’. Mutations at these sites can therefore render the binding ability of miRNA and thereby change the regulation as miRNA can be responsible for mRNA degradation and inhibition of translation.
*Result:*
Transcription: 1
Translation: 1
Protein: 0
*Example in literature:* 10.3390/cells8111465, 10.3390/genes8110296

**3'Flank:**

*Definition:* SO:“A sequence variant located 3' of a gene”
*Argument:* Possibly enhancers or isolators can be found here. Due to the change in transcription levels, the translational level would also be affected. This region is outside the coding region and will therefore not affect the protein.
*Result:*
Transcription: 1
Translation: (1)
Protein: 0

**RNA:**

*Definition:* SO:“A transcript variant of a non coding RNA gene”
*Argument:* Some non-coding RNA have the capability to inhibit translation or regulate degradation of mRNA. Non-coding RNA will not become a protein.
*Result:*
Transcription: 1
Translation: 1
Protein: 0
*Example in literature:* 10.3390/ijms20246249

**IGR:**

*Definition:* SO: “A sequence variant located in the intergenic region, between genes”
*Argument:* An intergenic region will not be part of a coding region. Possible regulatory regions could be enhancers, silencers or insulators not immediately up or

downstream of a gene. Due to the change in transcription levels, the translational level would also be affected. A MAF file (from TCGA) will only retain mutations in intergenic regions which are known to contain a regulatory region.

*Result:*

Transcription: 1
Translation: (1)
Protein: 0

*Example in literature:* 10.1007/978-1-60761-854-6_3
