## Supplementary Text 3 for "An Automatized Workflow to Study Mechanistic Indicators for Driver Gene Prediction with Moonlight"

**S3 Text. Methods used in case study applying Moonlight on Basal-like breast cancer.**

*Methods used in case study: Discovering driver genes in Basal-like breast cancer with Moonlight*

In order to demonstrate the application of the new functionalities in MoonlightR, a case study on Basal-like breast cancer was conducted on data collected from TCGA. Gene expression and mutational data from the TCGA-BRCA project were retrieved through the R/Bioconductor package TCGABiolinks v. 2.14.0., containing samples from *primary solid tumors* and *solid tissue normal*. Only female samples were retained. Both the gene expression and the mutational data were subtyped using TCGAbiolinks' function *PanCancerAtlas_subtypes()*, in which each sample is annotated with a molecular subtype. The breast cancer subtypes have been classified according to PAM50 signatures. We subsetted the gene expression and mutational data to include only the subtype Basal-like. We downloaded the mutational data as a MAF file and retained only the samples annotated with the somatic variant calling tool Mutect2, resulting in ~26,000 mutations across 171 patients. We filtered low quality variants from the somatic MAF file as those not fulfilling the following criteria: i) a minimum coverage of 30 from the tumor samples, ii) a minimum coverage of 10 from normal samples, iii) a variant read count of at least 3, and iv) a variant allele frequency (VAF) above 0.05. The reasoning behind these settings are explained below (see section "Quality analysis of MAF file"). This led to the retainment of around ~21,000 mutations across the 171 patients.
We processed the gene expression data in the following way using TCGAbiolinks: we removed outliers using the TCGAanalyze_Preprocessing function, normalized the data based on GC content and sequencing depth through the TCGAanalyze_Normalization function, and filtered out genes with low expression using TCGAanalyze_Filtering. The resulting gene expression matrix contained 17376 genes and 190 tumor samples. 170 patients were present in both the gene expression and mutation datasets, while one patient had only mutational data available and 20 patients only had transcriptome profiling data. Then, differential expression analyses (DEA) between the Basal-like subtype and normal samples were carried out using the limma-voom pipeline from the R/Bioconductor package limma (version 3.42.2) [1]. We downloaded clinical data from TCGA via TCGAbiolinks. The patients were primarily white and between the age of 40 and 70 when diagnosed.
Following the DEA, we executed the Moonlight pipeline. First, an enrichment analysis was performed with FEA, and two biological processes with opposing effects on cancer growth were chosen for the downstream pipeline: *proliferation of cells* and *apoptosis*. The two biological processes had an FDR of 3.9*10^-140 and 3.5*10^-92, respectively, and Moonlight Process Z-scores of 5.6 and -0.7, respectively. We performed GRN using the following parameters: kNearest = 3, nGenesPerm = 1000, and nBoot = 100. We conducted the remaining parts of Moonlight via the expert-based method, utilizing *apoptosis* and *proliferation of cells*. We identified the oncogenic mediators with *PRA()* with a threshold of zero. Finally, we ran the new function *DMA()*. The resulting tables from the DMA are used to select genes and mutations of interest. We focused this analysis on the 25 genes with the highest number of mutations and the genes containing driver mutations with transcriptional level of consequence in promoter regions. These genes are investigated in the literature and cross-referenced with other databases such as COSMIC and TRRUST. COSMIC includes established hallmarks of cancer for some driver genes, while TRRUST lists transcription factor relationships. The driver genes were also subject to an enrichment analysis with the R package EnrichR] using the

databases GO Molecular Function 2021, GO Biological Process 2021, and KEGG 2021 Human.
We compared Moonlight's driver gene predictions with GUST. For this, we downloaded GUST's predictions of driver genes in the TCGA-BRCA dataset from their webpage which comprised 285 driver genes divided into 269 TSGs and 16 OGs.

*Quality analysis of MAF file*

A Mutation Annotation Format (MAF) from TCGA has been annotated with a variant calling pipeline, i.e. Mutect2. However, when the variants' qualities are checked in the published MAF, some variants stick out. This is an overview and short analysis of the variants in the TCGA-BRCA Basal-like MAF. Firstly, it is highly noticeable that seven patients carry a large portion of the variants and are outliers on the 99-percentile (Panel A in S2 Fig). We will therefore also investigate whether these are significantly different in quality compared to the remaining patients' variants. These patients were either early stage or locally advanced, and none were metastatic.
There are only four metrics on which we can measure quality retained in the public MAF, which are normal depth, tumor depth, tumor reference read count and tumor variant read count, respectively. For some reads, the variant read count is as low as one, and for 2063 variants it is below 3. The depths were for more than 4000 variants below 30, and for 431 variants even below 10. It was also possible to calculate the variant allele frequency (VAF):

$$VAF = T_{alt\ count} \div t_{depth}$$

The depth in both tumor and normal is quite low, with an average for both between 80-120 reads, though a depth of at least x200 is recommended for somatic cells [2]. The VAF distribution is also trending to the lower end as the majority of the variants have VAF below 0.25.
After this analysis, we decided to run the case-study of Basal-like Breast Cancer from TCGA-BRCA, with some additional filtering of variants. We based the filtering on how well we could make the seven patients with a high number of mutations fit to the remaining average samples. When we filter on the tumor alteration read count to be at least five, the two patients: TCGA-AR-A0TU AND TCGA-AR-A0U0 are lowered to variant count of 206 and 275. We further sat a threshold of tumor depth of 30, normal depth of 10 and a VAF above 5%. This reduced the patient TCGA-A0SO to 368 variants.
After the filtering, four patients had still significantly higher mutation rates (Panel B in S2 Fig). These four patients were all white, aged between 55 and 81, all were alive at last follow up. Their tumor stages were between 1A-IIB, thereby not indicating that these patients are outliers in any other aera. It was decided that further analysis of these patients and their mutations were outside the scope of this paper.

**References**

1. Ritchie ME, Phipson B, Wu D, et al. Limma powers differential expression analyses for RNA-sequencing and microarray studies. Nucleic Acids Res 2015; 43:e47
2. Chen Z, Yuan Y, Chen X, et al. Systematic comparison of somatic variant calling performance among different sequencing depth and mutation frequency. Sci Rep 2020; 10:
