## Supplementary material for "An Automatized Workflow to Study Mechanistic Indicators for Driver Gene Prediction with Moonlight": Table S2

| **S2 Table. List of 286 Basal-like driver genes predicted by Moonlight divided into 87 OGs and 199 TSGs.** | | | | | |
|---|---|---|---|---|---|
| **OGs predicted by Moonlight** | | | | | |
| AHCY | AMPD1 | APC | BANK1 | BUB1 | BUB1B |
| CDC42BPB | CENPF | CHST15 | CLEC5A | CLEC9A | CPNE5 |
| DCAF13 | DERL3 | DLGAP5 | DOCK4 | DPH2 | EBNA1BP2 |
| ECM1 | EIF2AK1 | EMP1 | EMX2 | FAM13B | FAM162A |
| FANCI | FAT3 | FCRL2 | FCRL5 | GLRX2 | GTPBP4 |
| HERC1 | IL5RA | ILF3 | INCENP | ITPR1 | JAK1 |
| KCND2 | KIAA1109 | KIF2C | KIFC1 | KLF11 | KRTCAP2 |
| LGALS2 | MAD2L1 | MALSU1 | MCM2 | MFAP5 | MMS22L |
| MND1 | MPLKIP | MYO9A | NCBP1 | NFAT5 | ORC1 |
| ORC5 | PDLIM3 | PLEKHM3 | POLR2J | PRRC2B | PSMB4 |
| PTPRD | RAB8B | RACGAP1 | RAD54L | RPA3 | RPP38 |
| RUNX2 | SARAF | SEMA4D | SF3B4 | SGCD | SIGLEC6 |
| SLC12A3 | SLC12A4 | SULF1 | SYK | TCP1 | TCP11L2 |
| TENM3 | TOPBP1 | TOX | TREML2 | TROAP | UNC80 |
| VPS13C | VSTM4 | ZBTB8OS | | | |
| **TSGs predicted by Moonlight** | | | | | |
| A2M | ABCC9 | ACSM5 | ACVRL1 | ADAMDEC1 | ADAMTSL1 |
| ADAMTSL2 | ADCY4 | ADGRE1 | ADGRL4 | APBB1IP | APOL3 |
| ARHGAP25 | ARHGEF15 | ARL15 | ASPA | BAIAP2L1 | BCL6B |
| CACNA2D4 | CARD11 | CCDC178 | CCDC69 | CCM2L | CD200 |
| CD3G | CD48 | CD84 | CD86 | CDH5 | CERKL |
| CLNK | CMPK2 | COL14A1 | CPVL | CR1 | CRTAM |
| CSF1 | CST7 | CTSS | CX3CR1 | CXCL9 | CXCR6 |

| | | | | | |
|---|---|---|---|---|---|
| CYTIP | DDX58 | DEF6 | DOCK8 | EBF1 | EMCN |
| ERG | ETV7 | F5 | FAM124B | FBXL7 | FCGR1A |
| FCGR2A | FCGR3A | FCRL3 | FGD5 | FSTL1 | GMFG |
| GNAI2 | GRID1 | GZMA | HERC5 | HGF | HK3 |
| HLA-F | HSPA12B | HSPG2 | HTRA4 | IDO1 | IFFO1 |
| IFI44L | IFI6 | IFNG | IL12RB1 | IL21R | IL27 |
| IL2RA | IL2RB | INPP5D | ITGA1 | ITGAX | ITGB2 |
| KCNJ10 | KDR | KIF26B | KIR2DL4 | LAIR1 | LAMA2 |
| LAT2 | LCP1 | LDB2 | LILRA1 | LILRB3 | LIN7A |
| LTA | M1AP | MAP4K1 | MCM10 | MICAL2 | MICB |
| MIR155HG | MITF | MMP2 | MMRN2 | MNDA | MPEG1 |
| MS4A4E | MS4A7 | MX2 | NAPSB | NCF1 | NCF2 |
| NDUFS5 | NEFH | NIT2 | NLRC5 | NLRP3 | NR5A2 |
| NTHL1 | NUAK1 | OAS2 | OAS3 | OASL | PARP14 |
| PARP8 | PCDH17 | PCDHB12 | PCDHGB7 | PCOLCE | PDCD1 |
| PDGFB | PDGFRA | PEAR1 | PGR | PIK3R6 | PLAC9 |
| PRDM1 | PREX2 | PRKD1 | PSTPIP1 | PTPN7 | PTPRB |
| PTPRE | PTPRO | RASGRP4 | RELN | RUFY4 | SAMSN1 |
| SCARF1 | SELL | SEMA5A | SERPING1 | SH2D3C | SHE |
| SIGLEC10 | SIGLEC7 | SIGLEC9 | SIRPB1 | SIRPG | SLAMF1 |
| SLAMF6 | SLAMF7 | SLAMF8 | SLC15A3 | SLC17A9 | SLC24A4 |
| SLCO2A1 | SMG6 | SNX29 | SPARC | STAB1 | STX11 |
| SYNE1 | SYNE3 | TAL1 | TBC1D2B | TCF4 | TEK |
| THSD7A | TIGIT | TLL1 | TLR6 | TMEM132C | TMEM150B |
| TMEM204 | TNFRSF14 | TNFSF4 | TNIP3 | TPK1 | TRAT1 |
| TRIM14 | TTC24 | UBA7 | UBASH3A | VWF | WRNIP1 |
| XCL1 | | | | | |

Abbreviations: OGs, oncogenes; TSGs, tumor suppressor genes.
