## Supplementary material for "An Automatized Workflow to Study Mechanistic Indicators for Driver Gene Prediction with Moonlight": Table S3

**S3 Table. Comparison between Moonlight's predicted driver genes and driver genes reported in NCG.**

| **S3 Table.** Overlapping driver genes between Moonlight's predicted driver genes (TSG/OG) and driver genes reported in the NCG (TSG/OG). | | |
|---|---|---|
| **Moonlight's predicted driver type** | **NCG driver type** | **Overlapping genes between Moonlight's predicted driver type and the NCG database** |
| TSG | TSG | EBF1, NTHL1, PRDM1, PTPRB, TNFRSF1 |
| OG | OG | JAK1, SYK |
| OG | TSG | APC, BUB1B, PTPRD |
| TSG | OG | CARD11, ERG, KDR, MITF, PREX2, PDGFB, PDGFRA, TAL1 |
| Abbreviations: OG, oncogene; TSG, tumor suppressor gene; NCG, Network of Cancer Genes. | | |
