## Supplementary material for "An Automatized Workflow to Study Mechanistic Indicators for Driver Gene Prediction with Moonlight": Table S4

**S4 Table. Comparison between Moonlight's predicted driver genes, driver genes reported in NCG, and driver genes predicted by GUST.**

| **S4 Table.** Six genes which were predicted as either TSG or OG for breast cancer in both Moonlight, GUST, and NCG. | | | |
|---|---|---|---|
| **Moonlight predicted driver type** | **GUST predicted driver type** | **NCG driver type** | **Gene** |
| TSG | TSG | Candidate | CCDC178, PRKD1 |
| TSG | TSG | - | IFF01, HERC5 |
| OG | TSG | OG | JAK1 |
| OG | TSG | - | RUNX2 |
| Abbreviations: OG, oncogene; TSG, tumor suppressor gene; NCG, Network of Cancer Genes. | | | |
