## Supplementary figures and images for "An Automatized Workflow to Study Mechanistic Indicators for Driver Gene Prediction with Moonlight"

### Figure S1

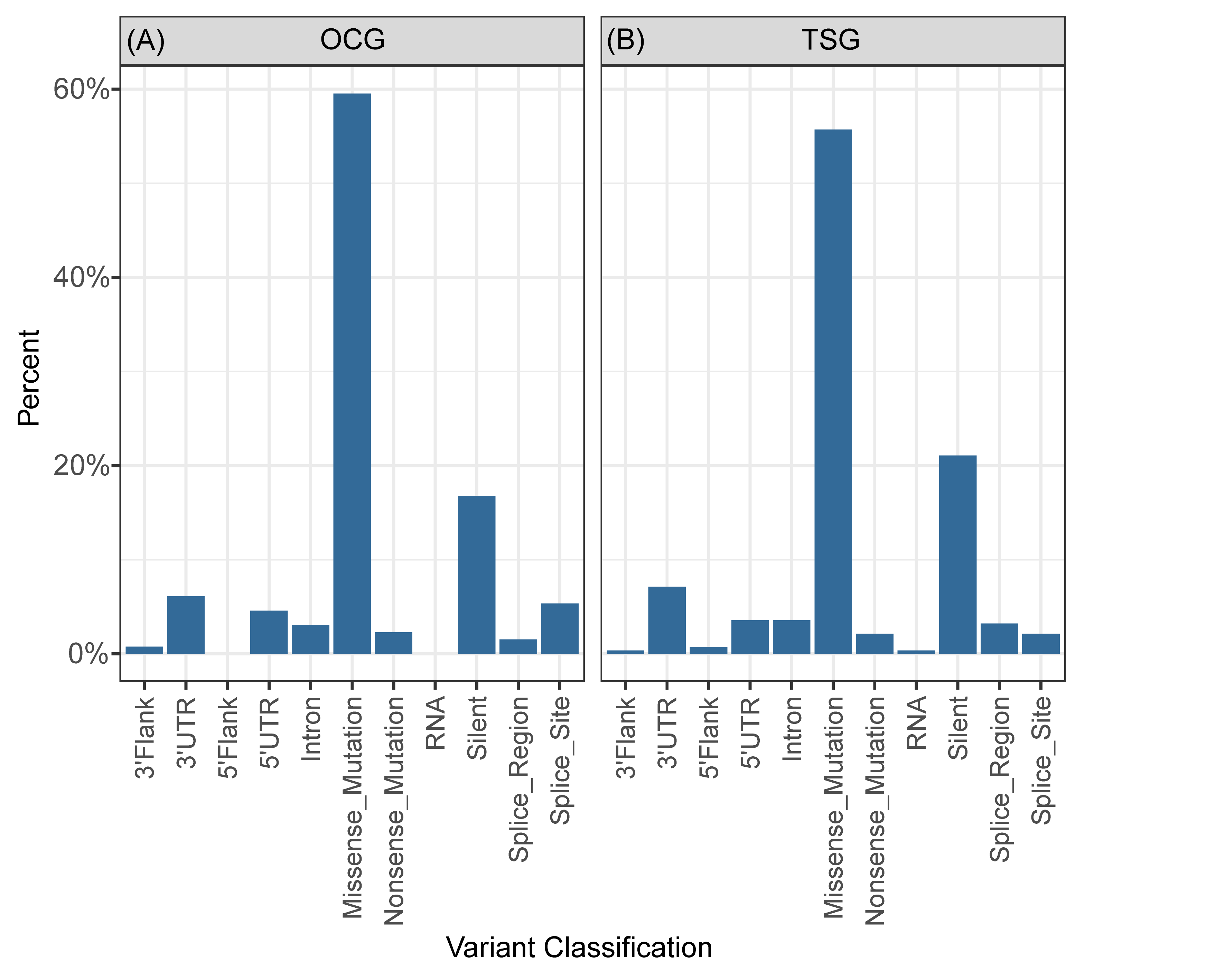

### Figure S2

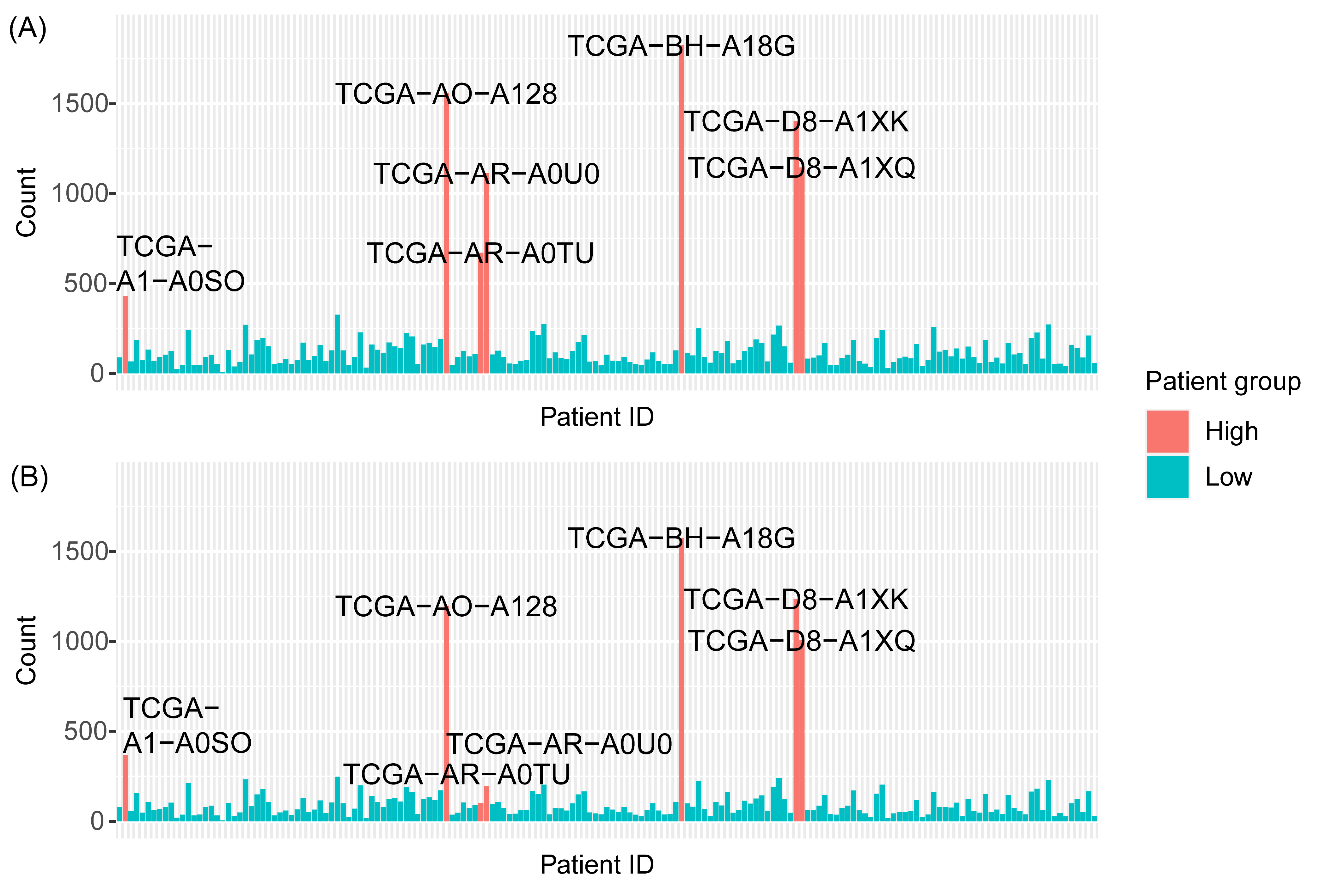
